# Design of a Computational Intelligence System for Detection of Multiple Sclerosis with Visual Evoked Potentials

**DOI:** 10.1101/2023.12.13.571427

**Authors:** Moussa Mohsenpourian, Amir Abolfazl Suratgar, Heidar Ali Talebi, Mahsa Arzani, Abdorreza Naser Moghadasi, Fariba Moghaddam, Seyed Matin Malakouti, Mohammad Bagher Menhaj

## Abstract

In this study, a new approach for modification of membership functions of a fuzzy inference system (FIS) is demonstrated, in order to serve as a pattern recognition tool for classification of patients diagnosed with multiple sclerosis (MS) from healthy controls (HC) using their visually evoked potential (VEP) recordings. The new approach utilizes Krill Herd (KH) optimization algorithm to modify parameters associated with membership functions of both inputs and outputs of an initial Sugeno-type FIS, while making sure that the error corresponding to training of the network is minimized.

This novel pattern recognition system is applied for classification of VEP signals in 11 MS patients and 11 HC’s. A feature extraction routine was performed on the VEP signals, and later substantial features were selected in an optimized feature subset selection scheme employing Ant Colony Optimization (ACO) and Simulated Annealing (SA) algorithms. This alone provided further information regarding clinical value of many previously unused VEP features as an aide for making the diagnosis. The newly designed computational intelligence system is shown to outperform popular classifiers (e.g., multilayer perceptron, support-vector machine, etc.) and was able to distinguish MS patients from HC’s with an overall accuracy of 90%.

## 1. Introduction

Our arsenal of immunomodulatory therapy options against the inflammatory aspect of multiple sclerosis (MS) has grown during the last ten years. However, it remains unclear which Disease-Modifying Treatment (DMT) is best for a given patient [1, 2, 3]. As a result, the decisions are the result of trial and error, and DMT changes and discontinuations are frequent occurrences [4,5, 6]. There is a dearth of information on the causes and timelines in clinical practice, particularly when it comes to recently diagnosed patients who have access to the most current DMTs.

Additionally, treating MS in more rural practices with limited access to neuroimmunological knowledge presents a challenge due to its increasingly complicated alternatives. Although MS is prevalent across Finland, there are notable regional variations, with MS being most infrequent in North Karelia, Finland’s most eastern region [7]. This is a rather remote area with a sparse population and vast distances. The North Karelia hospital district is not a participant in the national MS registry [8], which has demonstrated that, in comparison to other DMTs, the use of natalizumab, alemtuzumab, ocrelizumab, or rituximab as the first DMT was linked to a lower risk of 5-year disability progression and relapse. The research also found that, after a median of 2.4 years, 12.4% of the patients who had begun treatment with another DMT subsequently advanced to natalizumab, alemtuzumab, rituximab, or ocrelizumab [9]. Following the release of novel oral treatments, recent data from Finland also indicated a rise in DMT switches [10]. On the other hand, unlike Denmark and Sweden [4], no information is available about specific treatment plans or the causes of DMT discontinuation or switch in Finnish MS patients throughout the present treatment period.

In the present study, we search to find and define VEP components independently of amplitude that are also associated with MS. Furthermore, performing stochastic, temporal and spatial analysis on VEP recordings may yield useful information that may not be well understood in its original form [11,12,13,14]. Although there is currently no cure for MS, the FDA has authorized a number of medications for its management. Treatment for exacerbations, disease-modifying treatments (DMTs), and symptomatic medicines comprise the three categories of current therapy [15]. DMTs that address inflammatory immunopathology, for instance, might delay the onset of functional impairments but are unable to alleviate symptoms. As a result, creating efficient and unique treatment modalities is crucial [16]. Transcranial direct current stimulation (tDCS) in particular has gained popularity as a potential non-pharmacological therapeutic modality in recent years. Through the use of scalp electrodes, TDCS adjusts the resting membrane potential and provides low-current intensity, which can either increase or decrease the rate at which neurons fire. With opposing effects, the supplied current might be either positive or negative (anodal or cathodal stimulation, respectively): Excitatory post-synaptic potentials, which depolarize the neuronal membrane and alter cortical excitability, are often increased by anodal tDCS. Whereas the membrane becomes hyperpolarized and inhibited by cathodal tDCS [17]. Using highly repeatable and rater-independent methodologies to assess disease burden is one of the biggest unmet goals in MS research and clinical management. Low-contrast letter acuity measures, visual evoked potentials (VEP), optical coherence tomography (OCT), and efferent oculometrics have made significant strides in the measurement of visual dysfunction. However, most MS clinics do not routinely employ these tools, and they frequently need assistance from individuals with subspecialty training in neuroophthalmology [18]. The field of application for VEPs that use a diffuse flash stimulus, also known as flash-VEP or F-VEP, is rather limited when it comes to neurological pathologies affecting the visual pathways because these testing methods are less sensitive than P-VEP and produce incredibly variable responses in normal individuals. Additionally, P-VEPs enable the selection of features for the best stimulus in the clinical examination of the many visual system components, each of which may be triggered in a different way based on the image’s contrast, chromaticity, spatiality, and timing [19,20].

This information can offer a better diagnostic criterion in distinguishing normal subjects from subjects with neurological diseases, along with an index to indicate the progression of the diseases. Hence, it would be of great value to propose a pattern recognition system that can classify normal and MS subjects based on the unconventional features extracted from VEP signals.

The proposed computational intelligence system consists of five major elements: 1. preprocessing module for removing any possible artifacts in signals, 2. feature extraction module to calculate different components of the VEP signals, 3. feature subset selection module that selects effective components in diagnosis of MS, 4. classification module to distinguish signals based on their selected features, and 5. diagnosis module that indicates the class in which a signal belongs to. Figure 1 shows the block diagram of the proposed system.

**Figure 1:**
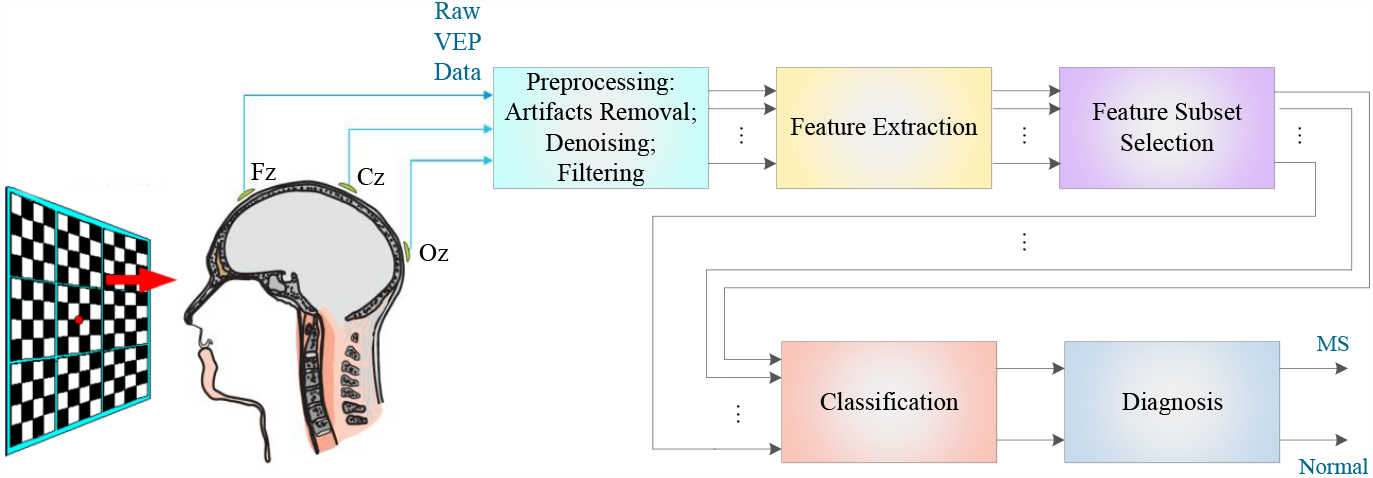
A block diagram of the implemented pattern recognition system.

## 2. VEP Recordings

### 2.1. Subjects

The baseline data is composed of 11 MS patients (median age 38.5 years; 90% female; median Expanded Disability Status Scale [20] (EDSS) 3.5, range 2–5.5; median disease duration 7.6 years, range 0.7–16 years) who were diagnosed with clinical isolated syndrome (n = 1; 9%), relapsing-remitting MS (n = 9; 81.8%), and secondary progressive MS (n = 1; 9%) based on the diagnostic criteria for multiple sclerosis [21]. The retrospective chart review was used in order to define history of optic neuritis (hON). Final diagnosis was made based on the clinical standard criteria: unilateral weakening or loss of vision over a period of hours or a few days, pain with eye movements, and declined perception of color. Ten patients (81%) had a positive history of ON. Also, ON was the first symptom in four patients (36%). Eleven subjects served as healthy controls (HC), having no remarkable personal history, accompanied by a normal brief neurological exam, while retaining a best corrected visual acuity of 0.8 or better (median age 39.5 years, 78% female).

### 2.2. Data Acquisition

A 4-chqnnel EMG system (Nihon Kohden MEB2200) at Sina Hospital was utilized in order to record Visual EPs. The recording, reference, and ground electrodes were placed with Oz, Fz, and Cz, respectively; with pre-auricular points used as landmarks. The impedance for electrodes was kept below 40 kΩ. Band-pass filter range for recording was set to 0.1–100 Hz, with the sampling frequency of 3 kHz. Pattern reversal VEPs were produced by full-field checkerboard stimulation independently applied to each eye, with compliance to international guidelines. Raw data went through a visual inspection, applied to a band-pass filter (1–30 Hz) and also averaged, while epochs with high amplitude artifacts were excluded. Recorded VEPs from MS patients and healthy controls are visualized in figures 2 and 3, respectively. Also, sample VEPs of two MS patients and two healthy controls are plotted in figures 4 and 5, respectively.

**Figure 2.**
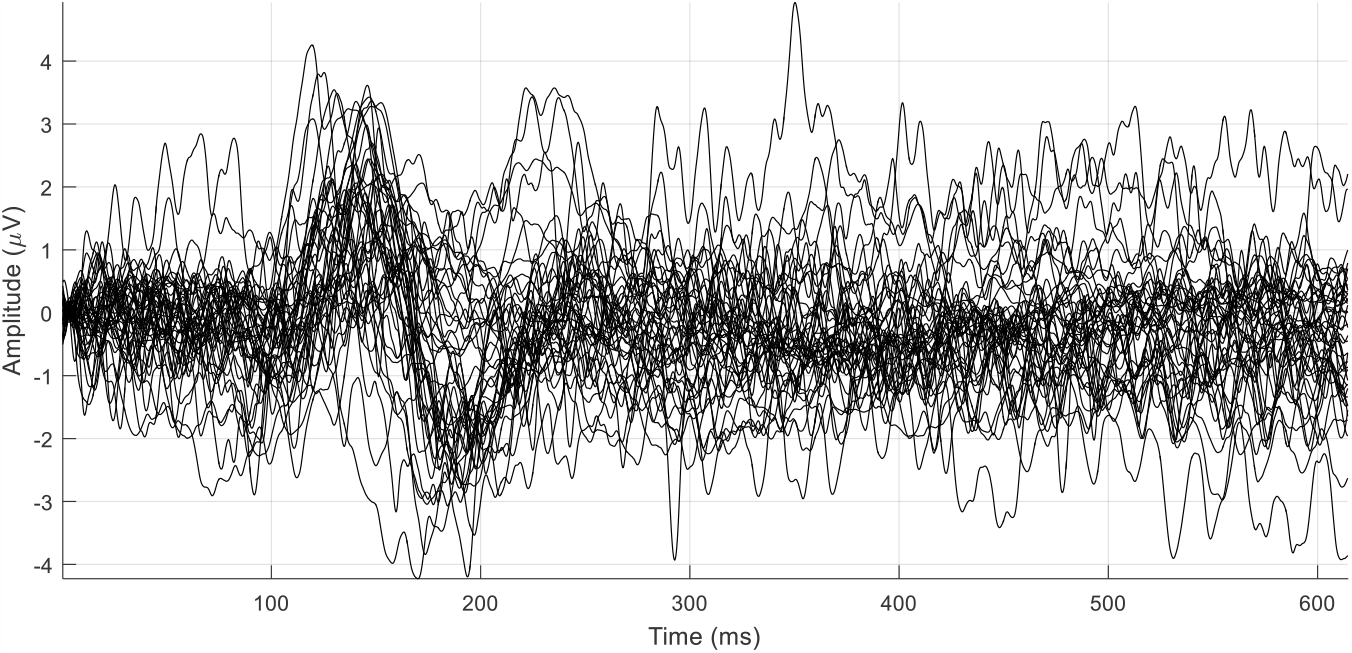
Pattern reversal VEP of all MS patients collected at Sina Hospital.

**Figure 3.**
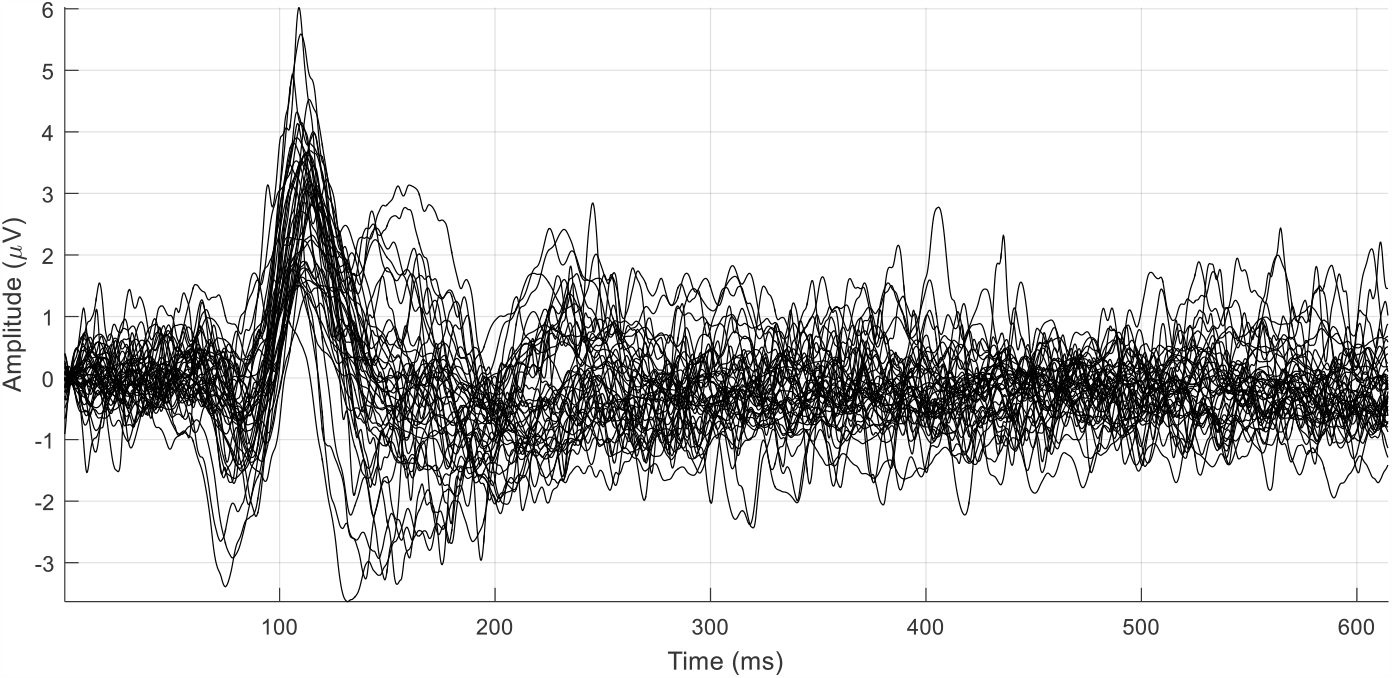
Pattern reversal VEP of all healthy subjects collected at the Sina Hospital.

**Figure 4.**
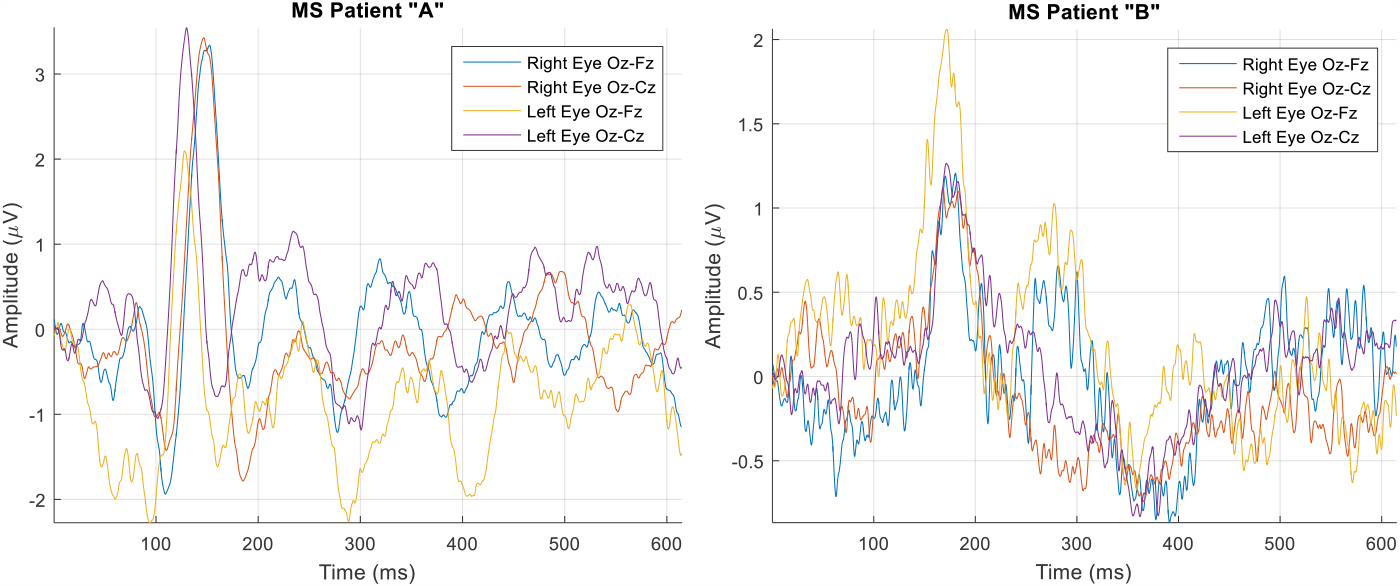
Two samples of pattern reversal VEP data collected from MS patients “A” and “B” at Sina Hospital.

**Figure 5.**
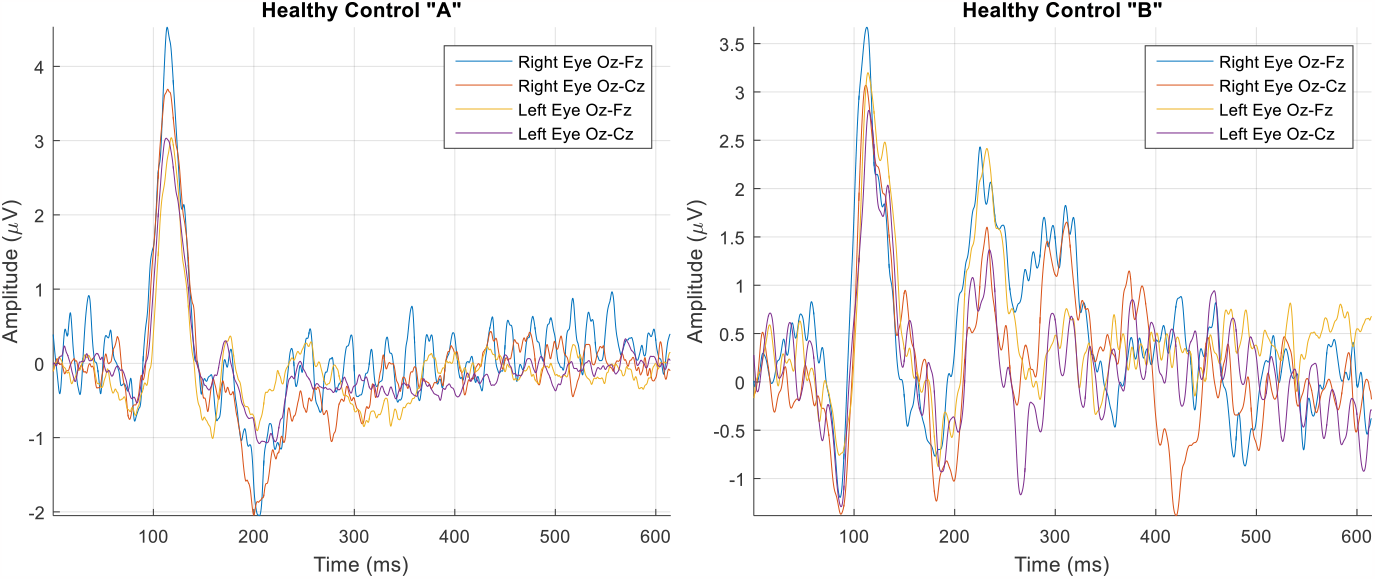
Two samples of pattern reversal VEP data collected from healthy controls “A” and “B” at Sina Hospital.

The VEP data for each subject consists of four arrays of waves, from which two were collected using the potential differences between Oz-Fz and Oz-Cz electrodes while only the subject’s right eye was open, and the other two were collected in a similar fashion while only the left eye was open. Accordingly, the data acquisition matrix is in the form of Equation 1.

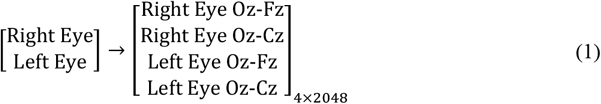

## 3. Extraction of the Features

### 3.1. Conventional Features

The aim of this section is to devise a method for extracting suitable features from the raw signal. The benefit of having a feature extraction module is to transform raw brain signals into a representation that can further simplify the classification. That is to say, feature extraction is an effort to remove as much noise and other redundant information from the input signals, while retaining information essential to distinguishing different classes of signals. Signal processing methods are used to extract feature vectors from the brain signals. This then allows for the comparison of the effect of various features on the performance of the detection system. Analysis of various time domain, frequency domain and time-frequency domain features resulted in the fact that features based on signal amplitude and time domain characteristics are more effective in revealing the P100 component [22]. These features are as follows:

Amplitude (AM, *C*_max_), which is the maximum signal value in [50,200] time interval:

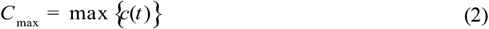

Positive value (PAV, *A*_*p*_ ), which is the sum of the positive values:

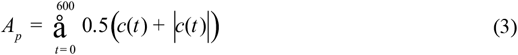

Latency 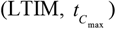, the P-VEP’s latency time, i.e. the time where the maximum signal value appears:

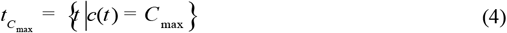

Where *c*(*t* ) is the P-VEP single trial during 0-600ms after stimulus and *C* _max_ is the maximum signal value in this time interval.

Negative area (NAV, *A*_*n*_ ), which is the sum of the negative signal values:

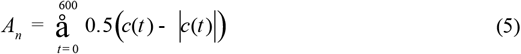

Peak-to-peak (PP, *pp* ):

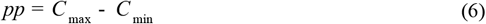

where *C* _max_ and *C* _min_ are the maximum and minimum signal values, respectively:

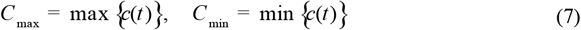

Peak of *N* 100 ( *P*_*N* 100_ ) the minimum signal value in [60, 190] time interval:

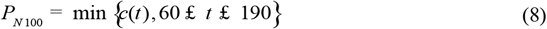

Latency of *N* 100 ( *P*_*N* 100_ ), the time where the *P*_*N* 100_ appears:

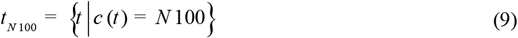

*P*1*N* 3, difference between the maximum signal value in [195, 550] time interval and the minimum signal value in [340, 500] time interval (corresponding to *P*100 amplitude and *N* 300 amplitude respectively).

*P*1*N* 1, difference between the maximum signal value in [195, 550] time interval and the minimum signal value in [70, 190] time interval (corresponding to *P*100 amplitude and *N* 100 amplitude respectively).

### 3.2. Wavelet Transform Features

In this section, the multi-scale wavelet transform features of VEP signals are calculated. These features include: Energy, Variance, Waveform Length, and Entropy. Mentioning that the VEP signal length is equal to 2048, since it was intended to divide the signal into two equal time windows, the window size was set to 1024, with a 1024 element incrementation designated for spacing of the windows. Decomposition level was assumed to be one. For a full tree at 1 level, 2 features are obtained. As it was decided to extract 4 types of features, the number of calculated features are 2 ’ 4 = 8 for each window, making for the total sum of 16 features for the two windows.

By inspection, it was observed that wavelet energy of the first window was quite the same among all VEP signals, and the wavelet variance was almost zero in all windows and all signals. Also, the wavelet entropy at Level 0 decomposition was almost equal to zero. Therefore, these 8 features were neglected, and the other eight were initially selected: Energy and Variance of first window at Level 1 decomposition; Energy and Variance of second window at Level 0 decomposition; and Energy, Variance, Waveform Length and Entropy of second window at Level 1 decomposition.

Overall, 17 features were extracted from each row of the data matrix. From these features, 9 were conventional signal components (i.e., amplitude, positive value, latency, negative area, peak-to-peak, peak of N100, latency of N100, P1N3, and P1N1), and the other 8 were those extracted from wavelet transformation of the signal (i.e., energy, variance, waveform length, and entropy). The feature extraction procedure is illustrated in Equation 10.

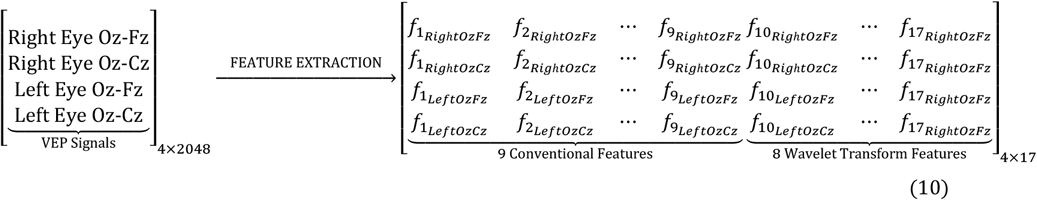

## 4. Feature Subset Selection

In this section, the objective is to select a suitable subset of extracted features in order to improve the classification of MS patients from healthy controls. For this purpose, dimension of the feature space needs to be reduced while making sure that the new feature subset results in a more efficient classification. This can be done through an optimization method called Direct Objective Optimization [23]. In this approach, first an ANN is created and trained using features extracted from the VEP.

Let **x** be a vector containing all (i.e., *n* _*f*_ ) features, 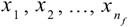, and **t** a vector containing all (i.e., *n* _*t*_ ) targets 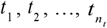:

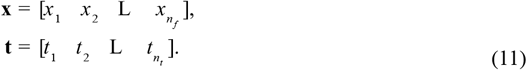

Now we assume **f** is a function taking feature values *x*_*i*_, *i* = 1, 2,K, *n*_*f*_ and returning its representation of target classes as *y*_*i*_, *i* = 1, 2,K, *n*_*t*_ in a vector 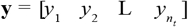:

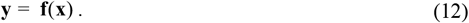

In order to avoid the complexity of the procedure **y** = **f**(**x**) (e.g., curse of dimensionality, Hughes effect, etc.), it is useful to reduce the number of features by eliminating those that are either redundant or less relevant. For this purpose, we assume 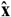 to be a vector containing a subset of original features *x*_*i*_, *i* = 1, 2,K, *n*_*f*_, namely 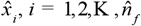 :

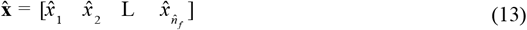

where obviously 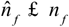 and 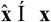 . Thus, a new function 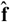 must be defined so that 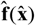 tracks **f**(**x**) with a negligible amount of error:

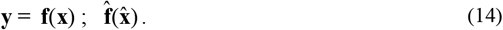

Now we define a tracking error **e** as follows:

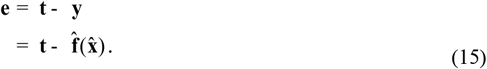

Hence, the mean squared error would be:

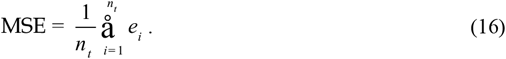

Now assuming that 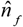 number of features needs to be selected, we can formulate a cost function as a weighted summation, in the following manner:

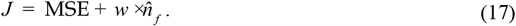

The weight *w* is defined to be proportional to the MSE (i.e., *w* μ MSE). Therefore, there is a coefficient *b* > 0 such that *w* = *b* ×MSE and 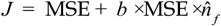 . Thus, the cost function can be rewritten as follows:

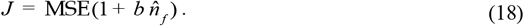

Here the goal is to define a scheme in which different subsets of 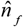 features compete with one another, and eventually the subset returning the best cost is selected. We define an array of decision variables 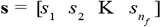 such that *s*_*i*_ Î {0,1}, *i* = 1, 2,K, *n*_*f*_ . If a decision variable, *s*_*i*_, *i* = 1, 2,K, *n*_*f*_, is equal to {1}, it would mean the corresponding feature, *x*_*i*_, *i* = 1, 2,K, *n*_*f*_, is selected. On the contrary, a feature *x*_*i*_ is not selected provided that its corresponding decision variable, *s*_*i*_, is set to be {0} . It is assumed that the capacity constraint (i.e., the number of features to be selected) is known and equal to 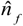. Therefore, the sum of {1} ‘s in the array **s** is also equal to 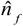 and we have:

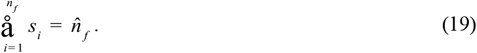

This can be viewed as a combinatorial optimization problem in which a permutation of features 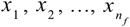 is created, and is ordered such that the first feature results in the best cost, while the last one causes the average cost to be the worst. Consequently, by choosing from the features that came first, a determined number of features (i.e., 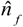 ) is selected and its corresponding cost is calculated. Two powerful discrete algorithms for solving combinatorial and permutative problems are Simulated Annealing (SA) [24], and Ant Colony Optimization (ACO) [25].

It was intended to select four most substantial VEP features from each eye (i.e., a total of eight features). In order to do so, both SA and ACO algorithms were implemented in MATLAB, according to the scheme described earlier. Both algorithms were run for 500 iterations. For SA algorithm, initial temperature was set to 10, with temperature reduction rate of 0.99, while having 20 sub-iterations within each iteration. For the ACO, number of ants was set to be 20, while setting the parameters as follows: initial pheromone, *t* _0_ = 1; pheromone exponential weight, *a* = 1 ; heuristic exponential weight, *b* = 1 ; and evaporation rate, *r* = 0.05 .

The SA was able to find an optimal solution with a cost equal to 1.7 ’ 10^-3^, and ACO’s best solution resulted in a cost equal to 3.6 ’ 10^-3^ . Therefore, the eight selected feature were the ones returned by the SA algorithm. These features were, from the Oz-Fz signals, Amplitude, Positive Value, and Latency for both eyes. Also, from the Oz-Cz signals of both eyes, the energies corresponding to the second window of wavelet transform at level 0 were selected.

## 5. Modification of Initial FIS with Krill Herd Optimization

Here, the objective is to modify an initial FIS structure by taking advantage of Krill Herd optimization algorithm (Figure 7). For this purpose, it is required to take the following steps:

**Figure 6.**
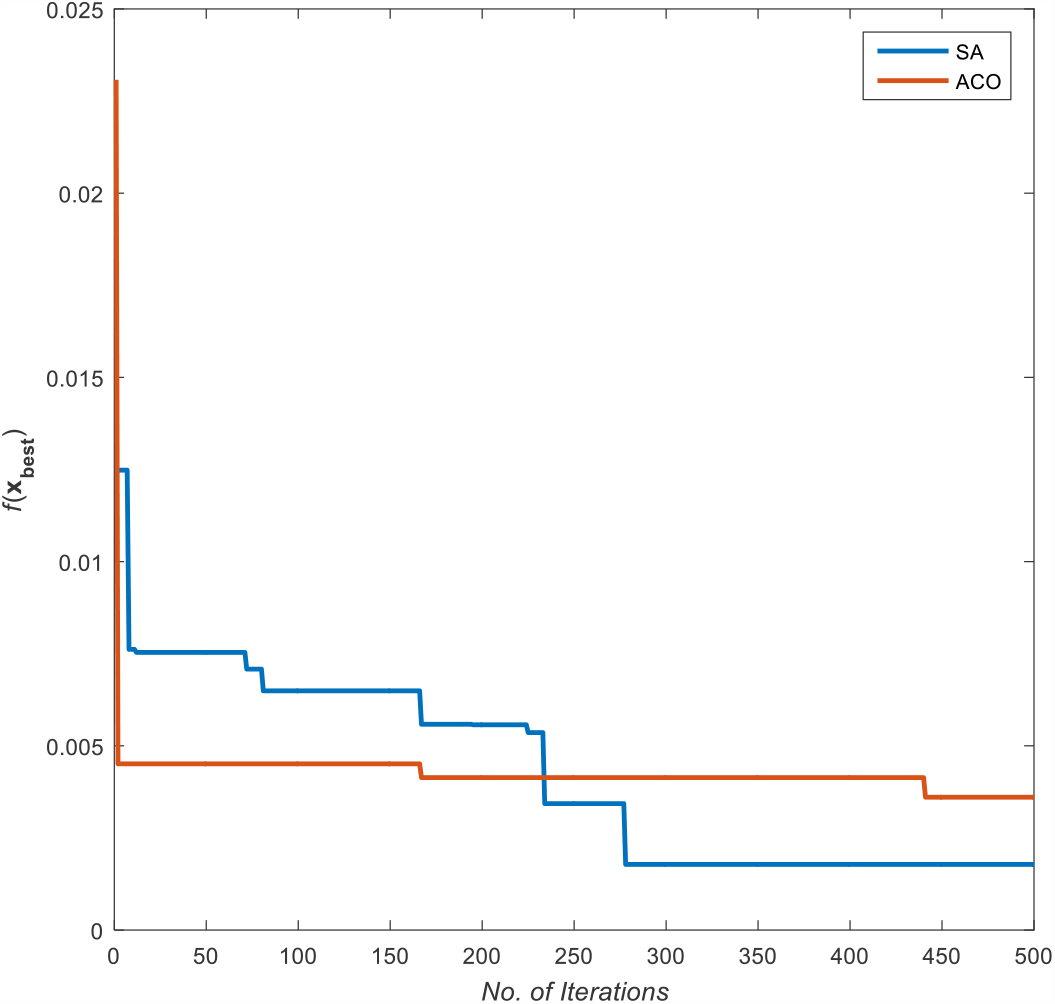
Best costs provided by ACO and SA optimization algorithms after 500 iterations.

**Figure 7:**
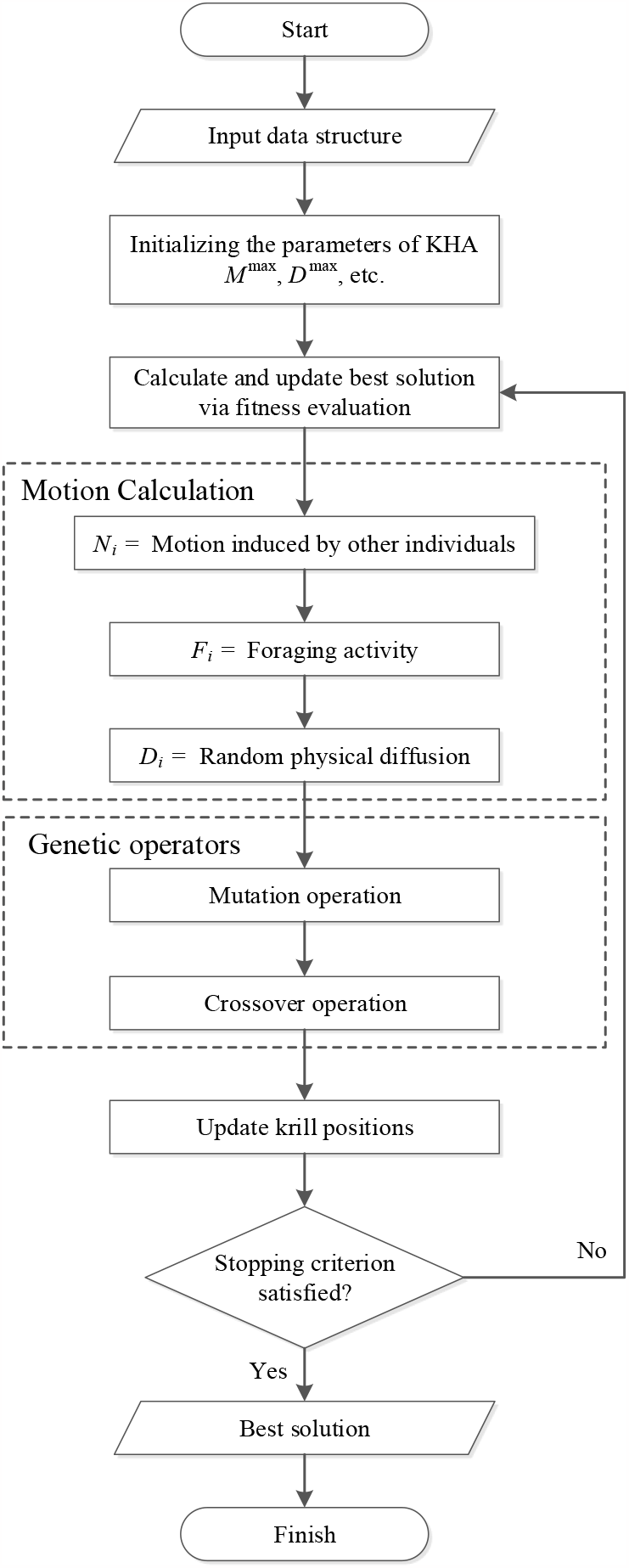
Simplified flowchart of the krill herd optimization algorithm.

1. Loading the training data
2. Creating an initial fuzzy inference system (FIS)
3. Modifying the values of FIS parameters according to modeling error, using the optimization algorithm
4. Returning the FIS with best values of parameters, as the final result

It is feasible to create an initial FIS structure using `genfis3` command in MATLAB. After applying the training data to the function `genfis3`, an FIS is generated using fuzzy c-means (FCM) clustering [26] by extracting a set of rules that models data behavior.

The rule extraction method first uses the function `fcm` in order to determine the number of rules and membership functions for the antecedents and consequents. In the case of our problem, more precisely a Sugeno-type fuzzy inference system [27], the input membership functions are Gaussian and have the following form:

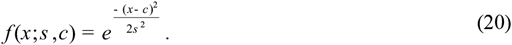

Moreover, membership functions for the outputs are linear and can be noted as follows:

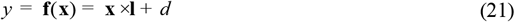

where *y* is a scalar, **f**(**x**) is a linear (i.e., affine) function, **x** is a 1-by-*m* vector, **l** is an *m* -by-1 vector, and *d* is a scalar.

Let 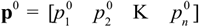 be a vector containing all membership function parameters (i.e., *s*_*i*_ ‘s and *c*_*i*_ ‘s for Gaussian, and vectors **l**_*j*_ and scalars *d* _*j*_ for linear membership functions). The goal is to find optimal values 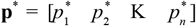 so that if substituted as modified membership function parameters, the training error is minimized. A cost function for the proposed optimization routine can be implemented as follows:

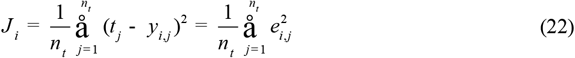

where *J*_*i*_, *i* = 1, 2,K, *n*_*it*_ is the cost value at iteration *i* ; *t* _*j*_, *j* = 1, 2,K, *n*_*t*_ is the *j* -th target; *y*_*i, j*_, *i* = 1, 2,K, *n*_*it*_, *j* = 1, 2,K, *n*_*t*_ is the value of *j* –th output of the network at iteration *i* ; and *e*_*i, j*_, *i* = 1, 2,K, *n*_*it*_, *j* = 1, 2,K, *n*_*t*_ is the training error of the *j* –th target at iteration *i* . The best resulted cost can be noted as *J* ^*^, which is also equal to the cost value at final iteration, 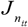 . The aim of optimization algorithm is to find a *w* belonging to the interval [-*M, M* ], *M* > 0 such that the equation

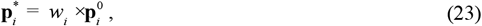

in each iteration results in the minimization of the cost, and hence, a more optimal set of parameters for fuzzy membership functions. *w*_*i*_ is calculated at the end of *i* -th iteration, and is multiplied by the initial values of parameters 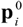 as a modification coefficient. **p**^*^ is calculated at the final iteration, and contains the best solution found by optimization algorithm to be set as parameters of membership functions. The optimization routine used for the purpose of our problem is a bio-inspired algorithm named Krill Herd [28].

## 6. Results and Discussion

Aforementioned classification methods were implemented using MATLAB. Since the dataset for MS patients and healthy controls was rather small (i.e., 11 MS and 11 HC), a random permutation of selected features was generated and presented as the input for pattern recognition systems. In each instance, 70% of the data was excluded for training, while the remaining 30% was used in order to test the validity of classification task. Classification procedures were run for 20 times, each for 100 epochs or iterations; and at the end all 20 training and testing results were averaged.

For the sake of comparison, three other more common classifiers were also employed: Multilayer Perceptron (MLP) [29], Support Vector Machine (SVM) [30] and Adaptive Neuro-Fuzzy Inference System (ANFIS) [31]. For MLP, two hidden layers of sizes 20 and 10 were implemented. The training and performance functions were set to Levenberg-Marquardt [32] and Cross-Entropy [33], respectively. Again, because of the relatively small dataset we were dealing with, the cross-validation option was neglected.

For the FIS and KH, the initial FIS was created with FCM clustering, and the number of clusters was set to 11. Exponent for the fuzzy partition matrix **U** was chosen as 2.0, while the minimum improvement in objective function between two consecutive iterations was selected to be 10^-6^ . For KH, number of runs was 3, population of the herd of krill was 30, and the crossover flag was set to 1. Also, *V*_*f*_, *D* ^max^, and *N* ^max^ were chosen to be 0.02, 0.005 and 0.01, respectively.

Regarding the SVM algorithm, data points were automatically centered at their mean, and scaled to have unit standard deviation, before training. Value of the box constraint **C** for the soft margin was set to 1, and the kernel cache limit was equal to 100. The kernel function was chosen to be linear, and the Karush-Kuhn-Tucker (KKT) [34] violation level was set to 0. The method used for finding the separating hyperplane was Sequential Minimal Optimization (SMO) [35], while the corresponding tolerance with which the KKT conditions are checked for the SMO training method was selected as 10^-3^ .

The corresponding confusion matrices for each classification is presented in the following tables.

As depicted in Table 5, the modification of FIS with KH optimization, compared to other three classification methods, led to the best classification results in accuracy, precision, sensitivity, specificity and diagnostic odds ratio (DOR) [36]. This shows that using KH algorithm for optimization of FIS training makes a powerful classifier that, in this case, can accurately and precisely separate the group of MS patients from healthy controls.

**Table 1.**
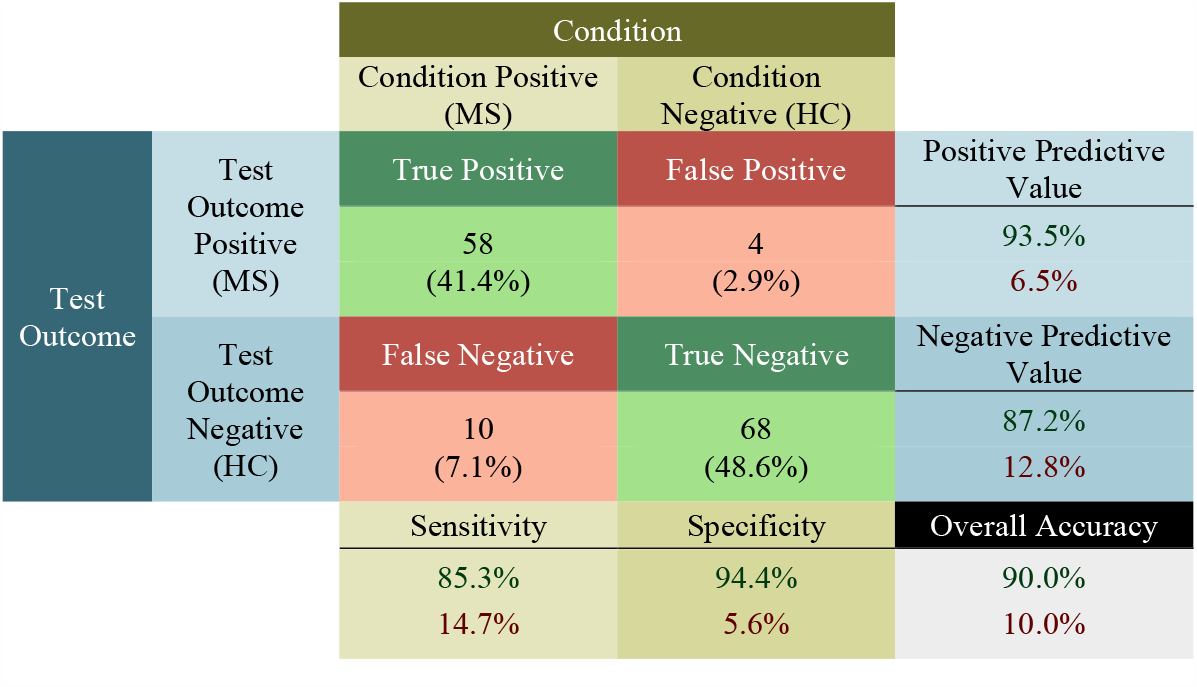
Confusion matrix for KH.

**Table 2.**
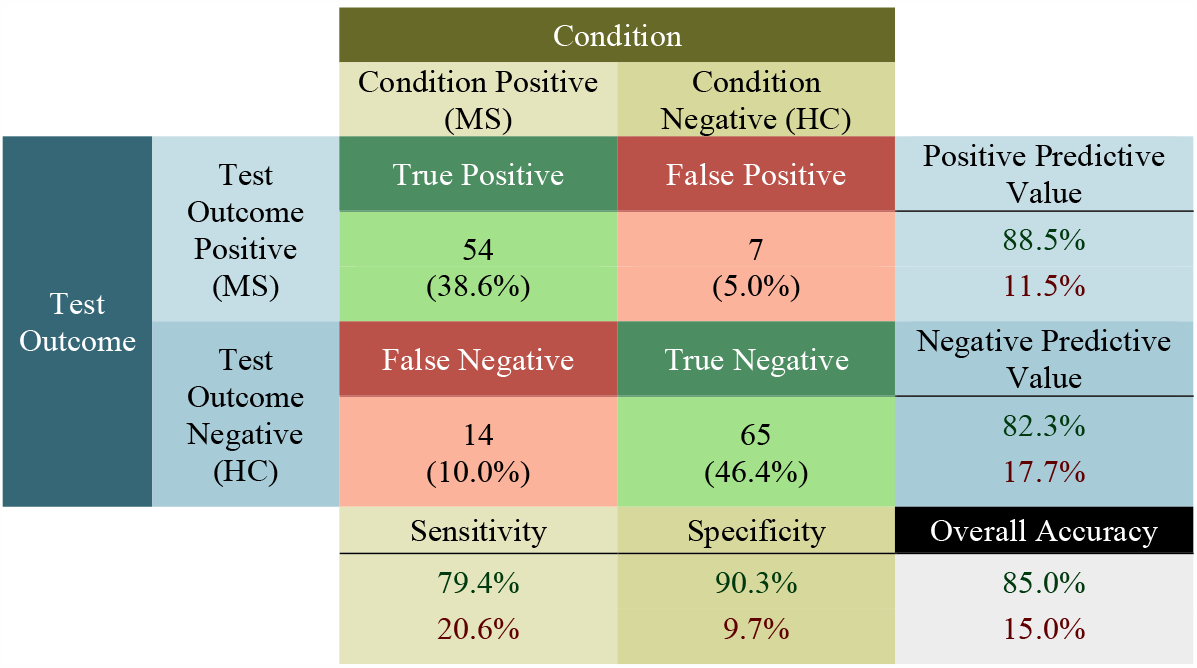
Confusion matrix for MLP.

**Table 3.**
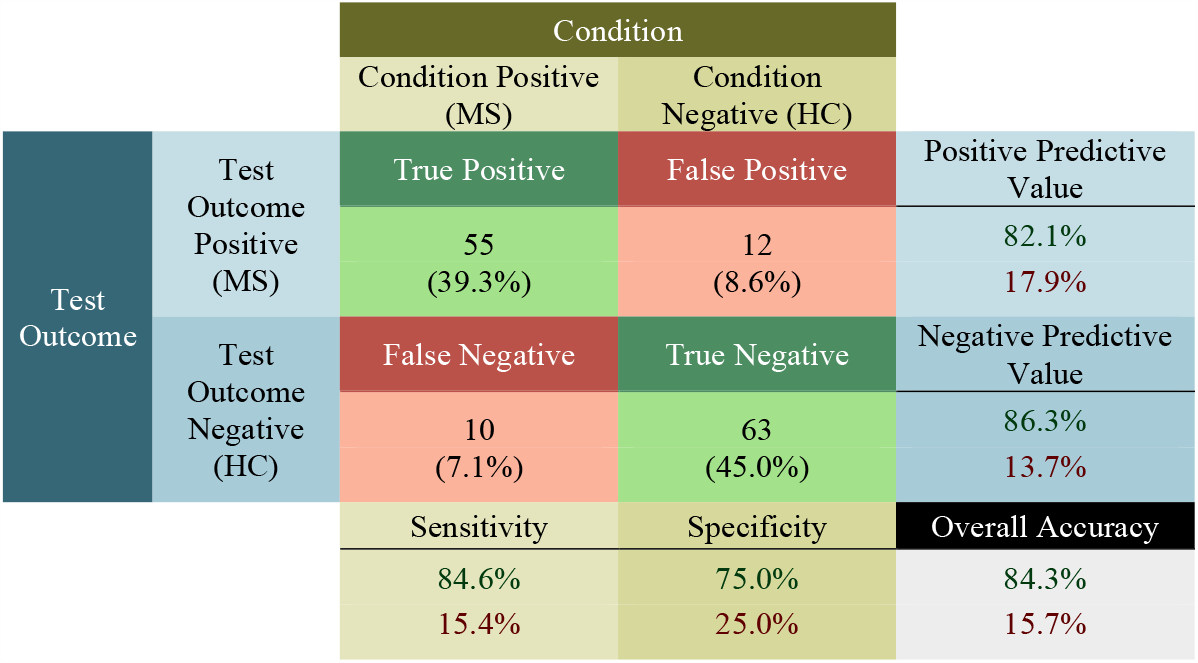
Confusion matrix for SVM.

**Table 4.**
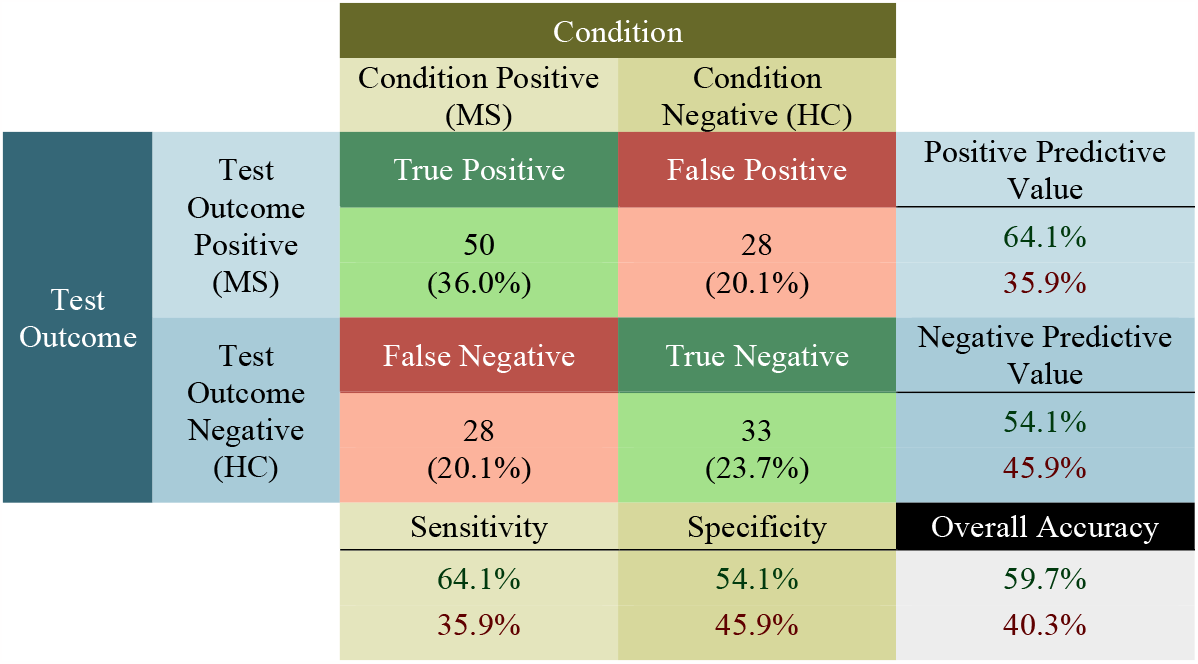
Confusion matrix for ANFIS.

**Table 5.**
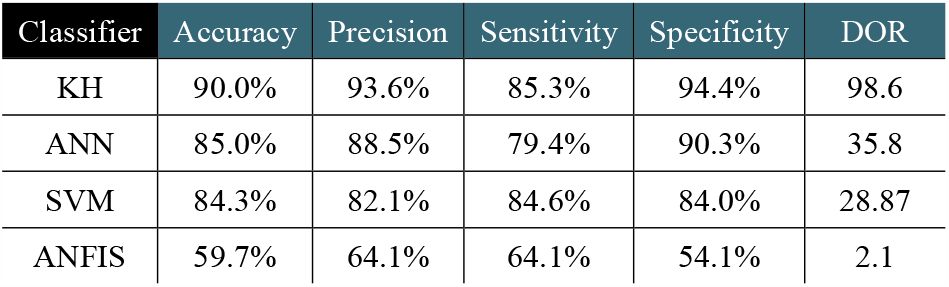
Comparison of the results of all classification methods.

| Classifier | Accuracy | Precision | Sensitivity | Specificity | DOR |
| --- | --- | --- | --- | --- | --- |
| KH | 90.0% | 93.6% | 85.3% | 94.4% | 98.6 |
| ANN | 85.0% | 88.5% | 79.4% | 90.3% | 35.8 |
| SVM | 84.3% | 82.1% | 84.6% | 84.0% | 28.87 |
| ANFIS | 59.7% | 64.1% | 64.1% | 54.1% | 2.1 |

## 7. Conclusions

This study demonstrated the use of KH optimization in modifying and training an FIS as a pattern recognition tool for classification of patients diagnosed with MS from healthy controls using VEP signals. As mentioned, the KH algorithm was utilized to adjust parameters associated with membership functions of both inputs and outputs of an initial Sugeno-type FIS, so that both training and testing errors were minimized. The application of this new pattern recognition system for classification of the VEP signals in 11 MS patients and 11 healthy controls was presented in Section 5.

As described earlier in Section 3, we also demonstrated the extraction of useful features from VEP signals. In Section 4, the most substantial features were selected in a feature subset selection scheme by taking advantage of two discrete optimization methods, ACO and SA. This selection of features provided further information regarding the value of many previously unused VEP features as an aide for making the diagnosis of MS.

Finally, the designed computational intelligence system was compared to other popular classification methods such as ANN, SVM and ANFIS (see Section 5). The new method was shown to outperform other classifiers and was able to distinguish MS patients from healthy controls with an overall accuracy of 90%.

In future, given the parameters associated with multiple sclerosis, such as MS subtype, disease modifying therapy (DMT), expanded disability status scale (EDSS) scores, and etc., a correlation analysis can be performed over the selected VEP features (see Section 4) to further understand the connection of VEP components with different aspects of the disease progression and prognosis. Topographic VEP (tVEP) also has been a hot topic in improvement of the MS diagnosis [37]. Topographic analysis of experimental recordings of VEPs may yield useful information that is not well understood in its original form. Such information may provide a good diagnostic criterion in differentiating normal subjects from subjects with neurological diseases, as well as an index of the progress of the diseases. Therefore, it would also be useful to apply considerable components of tVEP as inputs of a pattern recognition system, in order to see if the accuracy of classification can further be improved, hence achieving a less ambiguous computer-aided diagnosis of the disease.

## 8. Acknowledgements

Authors would like to thank all the MS patients as well as the staff at Sina Hospital who made this research possible. We also thank Vajiheh Amiraslani from the EEG and VEP team at Sina Hospital. for her assistance in collection of the data. A special thanks to Niloufar Mosharafian from the Computational Intelligence & Large Scale Systems Research Lab at Amirkabir University of Technology, for her gracious support and help throughout the study.

